# Genetically-encoded fluorescent biosensor for rapid detection of protein expression

**DOI:** 10.1101/2020.07.30.229633

**Authors:** Matthew G. Eason, Antonia T. Pandelieva, Marc M. Mayer, Safwat T. Khan, Hernan G. Garcia, Roberto A. Chica

## Abstract

Fluorescent proteins are widely used as fusion tags to detect protein expression *in vivo*. To become fluorescent, these proteins must undergo chromophore maturation, a slow process with a half-time of 5 to >30 min that causes delays in real-time detection of protein expression. Here, we engineer a genetically-encoded fluorescent biosensor to enable detection of protein expression within seconds in live cells. This sensor for transiently-expressed proteins (STEP) is based on a fully matured but dim green fluorescent protein in which pre-existing fluorescence increases 11-fold *in vivo* following the specific and rapid binding of a protein tag (*K*_d_ 120 nM, *k*_on_ 1.7 × 10^5^ M^−1^s^−1^). In live *E. coli* cells, our STEP biosensor enables detection of protein expression twice as fast as the use of standard fluorescent protein fusions. Our biosensor opens the door to the real-time study of short-timescale processes in research model animals with high spatiotemporal resolution.

---

*Aequorea victoria* green fluorescent protein (GFP) and its variants are widely used as quantitative reporters of gene expression to uncover the underpinnings of endogenous and synthetic circuits in contexts ranging from single cells in culture to whole animals.^*1-3*^ To become fluorescent, these proteins undergo chromophore maturation, an autogenic process that begins immediately following folding and involves successive steps of protein backbone cyclization, dehydration, and oxidation.^*4*^ The rate of chromophore maturation is highly dependent on temperature, pH, and oxygen concentration, which leads to large variations in half-times depending on experimental conditions.^*5*^ Under optimal conditions, maturation half-times for GFPs can be as low as 5 minutes in *E. coli*,^*5*^ but can increase to >30 min inside developmental model organisms such as frogs, zebrafish, and flies.^*6-8*^ These maturation half-times are too slow for quantitative detection of fast biological processes occurring within a few minutes, such as those involving transiently-expressed or fast-degrading proteins with half-lives of less than 5 minutes.^*9-11*^ As a result, accurate quantification of these proteins at a given point in time often requires *post hoc* mathematical transformations to correct delays in detection of protein expression caused by chromophore maturation.^*12-14*^

To minimize the delay between translation and detection of a protein of interest, biosensors that translocate a pre-expressed and fully-matured fluorescent protein from the cytosol to the nucleus following expression of a protein of interest have been developed.^*15, 16*^ However, the need for translocation prevents these biosensors from directly detecting proteins in the cytoplasm. Other biosensors use a repeating peptide fusion tag on the protein of interest to recruit multiple copies of a pre-expressed and fully matured cytosolic GFP, leading to the formation of large fluorescent aggregates that can be detected by fluorescence microscopy.^*17-19*^ While these biosensors enable real-time imaging of protein expression in individual cells, their large size (>1 MDa) can interfere with the physical properties of the protein of interest. Therefore, an ideal biosensor for the rapid detection of protein expression *in vivo* would not only minimize the delay between translation and detection of the protein of interest, but would also not require translocation of the fluorescent protein into a different subcellular compartment, or formation of large aggregates that may affect protein function.

Here, we create a genetically-encoded fluorescent biosensor to address these issues and enable the rapid detection of protein expression within live cells. We call our sensor STEP, for sensor for transiently-expressed proteins (Figure 1a). Inspired by the GCaMP family of biosensors that enable fast detection of Ca^2+^ dynamics,^*20*^ the STEP is based on a circularly permuted GFP (cpGFP) that can fold and mature independently of the protein of interest. In this cpGFP, the N- and C-termini are located in the middle of strand β7 of the β-barrel (Figure 1b), which creates a pore on the protein surface directly next to the chromophore phenolate moiety (Figure 1c). This pore exposes the chromophore to the solvent, resulting in quenched fluorescence (Figure 1a, OFF state).^*21*^ A peptide from the BH3 domain of the Bcl-2 family protein Bim^*22*^ is genetically fused to the N-terminus of cpGFP, creating a green fluorescent STEP (gSTEP). This Bim peptide enables specific binding of a protein tag (STEPtag) derived from another Bcl-2 family protein, Bcl-x_L_.^*23*^ Formation of the gSTEP/STEPtag complex causes a change to the electrostatic environment of the chromophore, restoring bright fluorescence (Figure 1a, ON state). By expressing gSTEP and allowing its chromophore to mature before expression of the STEPtagged protein of interest is initiated, the biosensor is ready to detect its target as it is expressed and folded, helping to eliminate delays in detection of protein expression caused by maturation.

**Figure 1.**
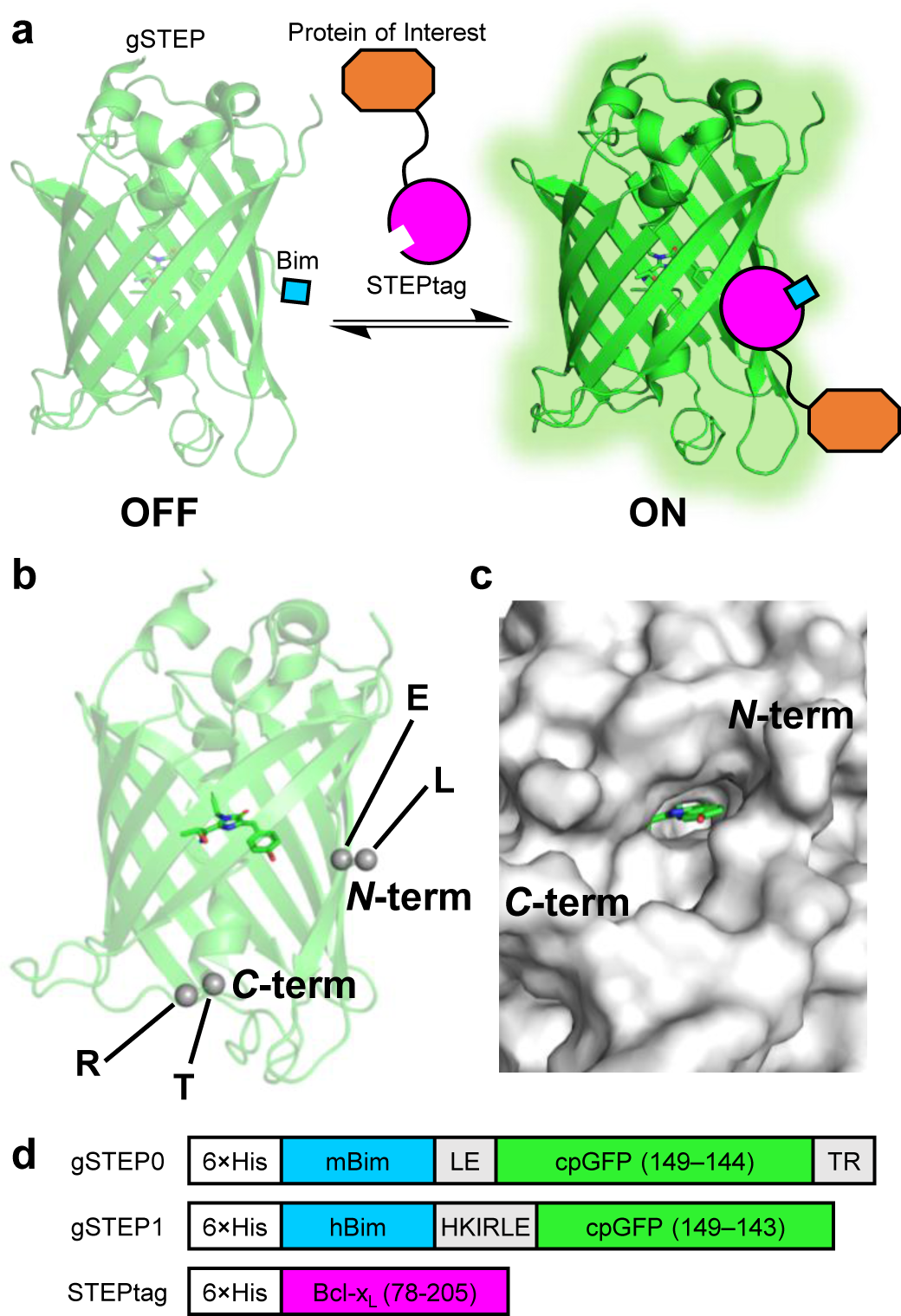
Sensor for transiently-expressed proteins (STEP). **a**, Cartoon representation of the STEP. A green fluorescent STEP (gSTEP) is expressed and allowed to mature before expression of a STEPtagged protein of interest (Not to scale). Prior to STEPtag binding to the Bim peptide, gSTEP is dimly fluorescent (OFF), while the bound gSTEP emits a strong fluorescence signal (ON). **b**, Crystal structure of the circularly-permuted GFP from the GCaMP3 genetically-encoded calcium indicator (PDB ID: 4IK8).^*40*^ The chromophore is shown as sticks, and residues forming the N- and C-terminal amino acid linkers are shown as grey spheres and identified by their one-letter code. **c**, Surface of the circularly-permuted GFP shows a pore on the barrel surface next to the chromophore phenolate moiety (green sticks). **d**, Schematic representation of gSTEP0, gSTEP1, and STEPtag. Linker sequences are shown in grey. Circularly-permuted GFP (cpGFP) is shown in green, and residues are numbered according to the sequence of *Aequorea victoria* GFP. Bcl-x_L_ is shown in magenta, and residues are numbered according to the UniProt sequence (Q07817). 6×His, mBim, and hBim indicate the histidine tag, mouse Bim, and human Bim peptides, respectively.

To create the first prototype of the sensor, gSTEP0, we fused the helical mouse Bim peptide (26 amino acids) to the cpGFP from the genetically-encoded calcium indicator GCaMP3,^*21*^ and retained the N- and C-terminal linkers on either side of the barrel pore (Leu-Glu and Thr-Arg, respectively), which have been shown to be important to the fluorescence response of these calcium sensors (Figure 1d, Supplementary Table 1).^*20*^ The STEPtag (15.5 kDa) was created by truncating the N- and C-termini of human Bcl-x_L_ (Figure 1d, Supplementary Table 1) to remove structural elements that are not essential for binding to Bim but can cause formation of a domain-swapped dimer,^*24, 25*^ and a hydrophobic membrane-anchor domain, respectively.^*26, 27*^ Addition of a saturating concentration of purified STEPtag to gSTEP0 resulted in an intensiometric fluorescence increase (ΔF/F_0_, calculated as (F_max_ − F_min_)/ F_min_) of 1.4 ± 0.1, with a dissociation constant (*K*_d_) of 250 ± 40 nM (Supplementary Figure 1, Table 1). Furthermore, control experiments confirmed that the fluorescence response of the biosensor was dependent on specific binding of the Bim peptide to the STEPtag (Supplementary Figure 1b,c).

**Table 1.** Properties of STEP variants

| Sensor | $\lambda_{\text{ex}}$<br>(nm)<sup>a</sup> | $\lambda_{\text{em}}$<br>(nm)<sup>a</sup> | $K_d$<sup>b</sup><br>(nM) | *In vitro*<br>$\Delta F/F_0$<sup>b</sup> | *In vivo*<br>$\Delta F/F_0$<sup>c</sup> | $k_{\text{on}}$<br>($\times 10^5 \text{ M}^{-1} \text{ s}^{-1}$)<sup>d</sup> | $k_{\text{off}}$<br>($\text{s}^{-1}$)<sup>e</sup> |
| --- | --- | --- | --- | --- | --- | --- | --- |
| gSTEP0 | 496 ± 1 | 513 ± 1 | 250 ± 40 | 1.4 ± 0.1 | N.D. | N.D. | N.D. |
| gSTEP1 | 504 ± 1 | 515 ± 1 | 120 ± 20 | 3.4 ± 0.4 | 11 ± 4 | 1.7 ± 0.2 | 0.020 ± 0.007 |

N.D. indicates not determined.
<sup>a</sup> n = 3, mean ± s.d. For comparison, excitation and emission wavelengths of EGFP are 488 and 507 nm, respectively.
<sup>b</sup> Measured in solution using purified gSTEP (75 nM) and STEPtag (up to 10 $\mu\text{M}$ ). For gSTEP0, n = 2 biological replicates, fit value ± 95% confidence interval. For gSTEP1, n = 6 biological replicates, fit value ± 95% confidence interval.
<sup>c</sup> Calculated from the average fluorescence of individual cells expressing both gSTEP1 and STEPtag, or expressing only gSTEP1 (see Figure 3a). Value represents the average of two biological replicates, and error is the standard deviation (n = 2, mean ± s.d.).
<sup>d</sup> Measured in solution using purified gSTEP1 (1 $\mu\text{M}$ ) and STEPtag (5 $\mu\text{M}$ ) (n = 3, fit value ± 95% confidence interval).
<sup>e</sup> Calculated from the $K_d$ and $k_{\text{on}}$ . Error represents the propagated 95% confidence interval.

Having established that gSTEP0 could be used to detect the presence of STEPtag *in vitro*, we next sought to improve the properties of our sensor. We began by truncating the C-terminus of gSTEP0 by removing the Thr-Arg linker (Figure 1d) as well as an additional 1 to 4 amino acids from cpGFP in order to increase the size of the pore on the barrel surface, which we hypothesized would improve ΔF/F_0_ by reducing background fluorescence through increased quenching in the unbound state. The best truncated mutant, gSTEP0-T1, had both the Thr-Arg linker and a single additional amino acid from cpGFP removed (Supplementary Table 1), and we found that it bound specifically to STEPtag with a *K*_d_ of 210 ± 80 nM and a ΔF/F_0_ of 2.1 ± 0.4 (Supplementary Figure 2, Supplementary Table 2). Control experiments with this improved variant confirmed that fusion of STEPtag using a 10-amino acid linker to either the N- or C-terminus of a protein of interest does not substantially affect biosensor response or binding affinity (Supplementary Figure 3).

**Table 2.** Time required to reach a specified level of fluorescence following induction of protein expression in live *E. coli* cells

| Fluorescent Reporter <sup>a</sup> | Time to reach X standard deviations above initial fluorescence intensity (min) <sup>b</sup> |  |  |
| --- | --- | --- | --- |
|  | X = 1 | X = 2 | X = 3 |
| gSTEP1 | 0.63 ± 0.03 | 1.09 ± 0.06 | 1.6 ± 0.2 |
| EGFP | 1.21 ± 0.09 | 3 ± 2 | 4 ± 1 |
| sfGFP | 1.1 ± 0.4 | 2.1 ± 0.6 | 2.9 ± 0.4 |

<sup>a</sup> gSTEP1 refers to cells expressing both gSTEP1 and STEPtag. EGFP and sfGFP refer to cells expressing only EGFP or sfGFP. STEPtag, EGFP, and sfGFP expression is under control of the araBAD promoter, and can be induced using arabinose. gSTEP1 is constitutively expressed.
<sup>b</sup> Fluorescence of the bacterial cell population was measured for 10 minutes before induction of STEPtag, EGFP, or sfGFP expression using 0.45% arabinose, and this baseline signal was used to calculate the standard deviation serving as detection threshold (n = 2 biological replicates, mean ± s.d.).

**Figure 2.**
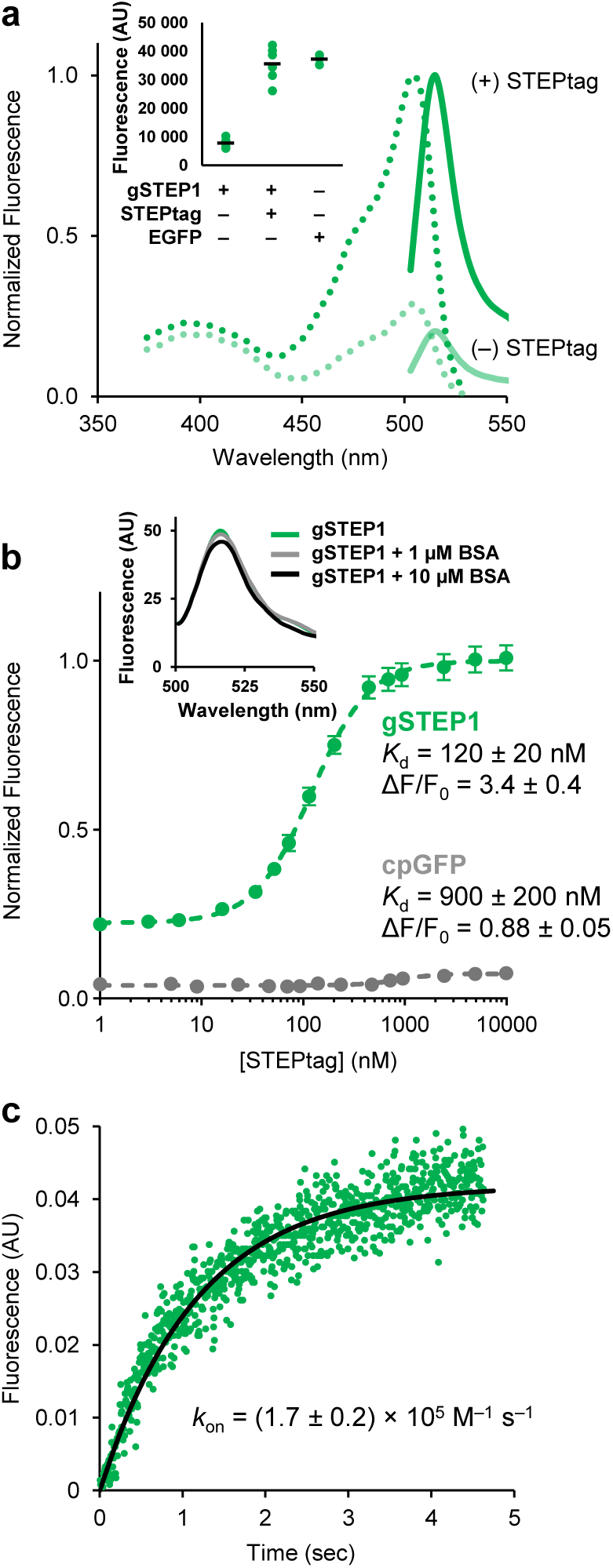
*In vitro* characterization of gSTEP1. All assays were performed in 20 mM sodium phosphate buffer containing 50 mM NaCl (pH 7.4). **a**, Normalized excitation (λ_em_ = 550 nm, dashed line) and emission (λ_ex_ = 485 nm, full line) spectra of gSTEP1 (75 nM) in the presence or absence of saturating STEPtag (10 µM). Inset shows the fluorescence intensity at 515 nm (λ_ex_ = 485 nm) of six biological replicates of gSTEP1, in the presence or absence of saturating STEPtag, compared to three technical replicates of 75 nM EGFP. Mean values are shown as black lines. **b**, Binding curves of 75 nM gSTEP1 (green) or cpGFP (grey) with STEPtag. Fluorescence is normalized to the maximum intensity observed for gSTEP1. Dashed lines represent fits of the Hill equation to the data (Hill coefficients of 1.5 or 2.2 for gSTEP1 or cpGFP, respectively). For the gSTEP1 binding curve, data points represent mean ± SEM of six biological replicates. For cpGFP, data points represent mean of three technical replicates. *K*_d_ and ΔF/F_0_ values were obtained from the fit and indicated with the 95% confidence interval around the fit values. Inset shows emission spectra (λ_ex_ = 485 nm) of 75 nM gSTEP1 in the presence of 0, 1, or 10 µM bovine serum albumin (BSA). **c**, Rapid-mixing stopped-flow binding kinetics of gSTEP1 mixed with saturating STEPtag. The black line represents a fit of the integrated rate equation to the data (Methods).

**Figure 3.**
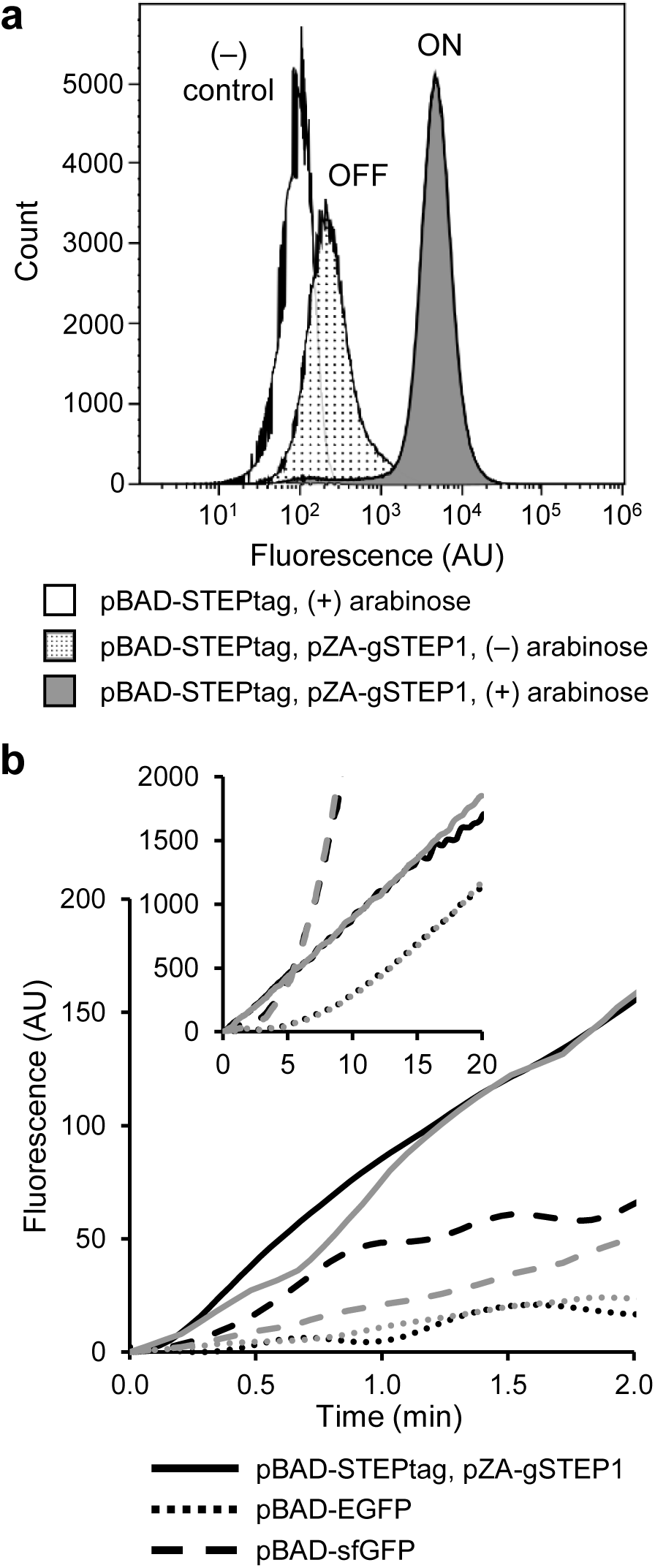
gSTEP1 enables rapid detection of protein expression in live bacterial cells. **a**, Flow cytometry histograms of gSTEP1 fluorescence in live *E. coli* cells expressing only STEPtag (negative control), gSTEP1 (OFF state), or both (ON state). The pZA vector constitutively expresses gSTEP1, while STEPtag expression from the pBAD vector is induced using 0.2% arabinose. **b**, Time course of protein expression in live *E. coli*. Cells were grown to the end of the exponential growth phase (OD600 = 1.1), then fluorescence was measured immediately after pBAD vectors containing either STEPtag (for cells constitutively expressing gSTEP1), EGFP, or sfGFP were induced with 0.45% arabinose. Two biological replicates (black and grey) are shown, each blanked by the fluorescence signal at 0 min, and smoothed by three passes through a seven-point moving average filter. Inset shows the 20-min time course. No fluorescence increase was observed by addition of arabinose to cells containing pZA-gSTEP1 and empty pBAD (Supplementary Figure 4).

Next, we replaced the mouse Bim peptide of gSTEP0-T1 with the human homolog or a range of synthetic variants displaying tight binding to Bcl-x_L_,^*28*^ which we hypothesized would enhance binding affinity to the STEPtag. Of these, the human Bim peptide performed the best (*K*_d_ = 170 ± 40 nM, ΔF/F_0_ = 3.3 ± 0.6, Supplementary Table 2). In parallel, we tested various linker lengths (1 to 5 amino acids) between the original mouse Bim peptide and cpGFP in gSTEP0-T1 to allow alternate binding poses of the STEPtag on the gSTEP surface upon formation of the complex. We hypothesized that changing the relative orientation of the binding partners could enhance binding affinity or ΔF/F_0_ by allowing more favourable non-covalent interactions between these molecules or causing a larger change to the electrostatic environment of the chromophore upon binding, respectively. We found that addition of a four-amino acid linker (gSTEP0-T1-L4) improved the binding affinity but not ΔF/F_0_ relative to gSTEP0-T1 (Supplementary Table 2). Interestingly, replacement of the mouse Bim peptide in gSTEP0-T1-L4 by its human homolog yielded a worse *K*_d_ and ΔF/F_0_ even though human Bim performed better than mouse Bim in gSTEP0-T1. Therefore, as a final step, we performed combinatorial saturation mutagenesis of the four-amino acid linker introduced between human Bim and cpGFP in gSTEP0-T1-L4, and screened the resulting library for improved brightness and ΔF/F_0_ using fluorescence-activated cell sorting and microplate-based binding assays, respectively (Methods). This yielded our final improved variant, gSTEP1 (Figure 1d, Table 1, Supplementary Table 1), which displays a ΔF/F_0_ of 3.4 ± 0.4, equivalent to that of the original GCaMP (ΔF/F_0_ = 3.5),^*20*^ and is as bright as the enhanced GFP (EGFP) from *Aequorea victoria*^*29*^ when fully bound to STEPtag (Figure 2a). gSTEP1 binds specifically (Figure 2b) and rapidly (Figure 2c) to STEPtag, with a *K*_d_ of 120 ± 20 nM and a binding rate constant (*k*_on_ = 1.7 ± 0.2 × 10^5^ M^−1^s^−1^) that is comparable to that of peptide antigen binding by antibodies.^*30*^

Next, we evaluated whether gSTEP1 could be used to detect STEPtag expression in live *E. coli* cells, which we selected as a case study given the fast GFP maturation rate in this organism.^*5*^ To do so, we prepared an *E. coli* strain that constitutively expresses a low basal concentration of gSTEP1 and in which STEPtag expression can be induced by the addition of arabinose (Methods). In flow cytometry experiments, we observed that cells constitutively expressing gSTEP1 and overexpressing STEPtag were considerably brighter than those that do not express the binding partner (Figure 3a), with little overlap between the fluorescence distributions of the two cell populations. Under these conditions, the mean fluorescence of the cellular population in the ON state was an order of magnitude higher than that of the cellular population in the OFF state, resulting in a ΔF/F_0_ of 11 ± 4 (Table 1). Taken together, these results demonstrate that the fluorescence difference of gSTEP1 in the ON and OFF states is sufficient to distinguish individual bacterial cells that express STEPtag from those that do not.

Having demonstrated that gSTEP1 could be used to detect the STEPtag in live *E. coli* cells at the steady-state, we evaluated the ability of the biosensor to report on STEPtag concentration dynamics. To do so, we cultured the cells constitutively expressing gSTEP1 until they reached the exponential growth phase, and then induced expression of STEPtag by adding arabinose. We observed an immediate fluorescence increase (Figure 3b), and the signal continued to increase linearly for 20 min. To determine how long it takes for protein expression to be detected by our biosensor, we measured the baseline fluorescence of these cells prior to induction of STEPtag expression (Supplementary Figure 4), and used the noise in this baseline data to set detection thresholds above the signal at time of induction (t = 0 min). The standard deviation was used to quantify the noise, such that the thresholds of 1, 2 and 3 standard deviations above the signal at t = 0 min represent increasing levels of confidence that the increase in fluorescence is due to the fluorescent reporter (Table 2). For cells expressing both gSTEP1 and STEPtag, the threshold of 3 standard deviations of the baseline above the signal at 0 min was reached in 1.6 ± 0.2 min. By contrast, when we induced expression of EGFP (maturation half-time = 25 min^*29*^) using the same promoter in cells containing only the EGFP expression vector, it took 4 ± 1 min for it to reach the same threshold, over twice as long as for gSTEP1. Of note, the rate of fluorescence increase for EGFP accelerated with time, reaching a steady state after approximately 10 minutes under these conditions. Presence of this lag phase is consistent with slower oxidation than folding/cyclization/dehydration during GFP chromophore maturation.^*31*^ In the first 5 minutes following induction of protein expression, gSTEP1 provided 6-to 10-fold higher fluorescence signal than EGFP, and this signal remained higher for approximately 30 minutes (Supplementary Figure 4). We also tested Superfolder GFP (sfGFP), which folds and matures faster than EGFP (maturation half-time = 13.6 min^*32*^). Expression of sfGFP using the same promoter also resulted in a lag phase, albeit shorter than the one observed for EGFP (approximately 5 minutes to reach steady-state), and yielded a fluorescence intensity increase of 3 standard deviations above the initial signal in 2.9 ± 0.4 minutes (Table 2). These results demonstrate that gSTEP1 enables faster detection of protein expression in live *E. coli* cells than the use of traditional GFP reporters, which should increase the temporal resolution of experiments aiming to detect transiently-expressed proteins or other fast biological processes.

Compared with other genetically-encoded fluorescent biosensors used to track protein expression in real-time, gSTEP1 has the benefits of not requiring the use of protein translocation^*15, 16*^ or formation of large protein aggregates,^*18*^ which should cause minimal perturbation to the subcellular localization and physical properties of the protein of interest. In the course of this work, a protein biosensor operating on a similar principle to the STEP was published.^*33*^ This sensor, called Flashbody, is based on a cpGFP that is inserted between heavy and light chain fragments from the variable region of an antibody, which together bind specifically to a 7-amino acid peptide tag fused to a protein of interest. Like gSTEP1, the Flashbody has the benefits of not requiring translocation or formation of large aggregates, and the response of the two biosensors to their respective binding partner is similar (ΔF/F_0_ ≈ 3). However, gSTEP1 displays tighter binding (*K*_d_ of 120 nM for gSTEP1 vs. 423 nM for the Flashbody), which could allow detection of proteins present at lower concentrations than the Flashbody limit of detection, and binds to its partner with a rate constant two orders of magnitude higher than that of the Flashbody (*k*_on_ of 1.7 × 10^5^ M^−1^s^−1^ for gSTEP1 vs. 3.38 × 10^3^ M^−1^ s^−1^ for Flashbody).^*33*^ Taken together, these advantages of gSTEP1 make it a useful alternative to other biosensors for the rapid detection of protein expression *in vivo* and in real time.

In conclusion, we have developed a genetically-encoded fluorescent biosensor to rapidly detect protein expression within live cells. Because it is based on a circularly permuted GFP, our sensor should be applicable for use in research model animals. However, for some applications, it may be necessary to further improve the biosensor’s dynamic range and sensitivity. This could be achieved by replacing the Bim/STEPtag pair by alternate binding partners, and optimizing the fluorescence response by random mutagenesis followed by rounds of fluorescence-activated cell sorting using the pZA-gSTEP1/pBAD-STEPtag strain developed here to allow modulation of the STEPtag concentration. Alternate colors should also be possible via the use of circularly permuted yellow^*34*^ or red^*35*^ fluorescent proteins. We expect that the engineering of a color palette of orthogonal STEP biosensors will enable multiplexing for more complex imaging experiments, opening the door to the *in vivo* visualization of protein concentration dynamics in real time and at unprecedented spatiotemporal resolution.

## Acknowledgements

R.A.C. acknowledges grants from the Canada Foundation for Innovation (26503) and the Human Frontier Science Program (RGP0041). H.G.G. was supported by the Burroughs Wellcome Fund Career Award at the Scientific Interface, the Sloan Research Foundation, the Human Frontiers Science Program (RGP0041), the Searle Scholars Program, the Shurl & Kay Curci Foundation, the Hellman Foundation, the NIH Director’s New Innovator Award (DP2 OD024541-01), and an NSF CAREER Award (1652236). M.G.E. and M.M.M. are the recipients of Ontario Graduate Scholarships and postgraduate scholarships from the Natural Sciences and Engineering Research Council of Canada. The authors thank Shahrokh Ghobadloo for assistance with the fluorescence-activated cell sorting and flow cytometry experiments, Jeffrey W. Keillor for use of the rapid-mixing stopped-flow spectrophotometer, James A. Davey for providing the EGFP gene, and Stephen L. Mayo for providing the *Thermoascus aurantiacus* xylanase 10A expression vector.

## Author contributions

R.A.C. and H.G.G. conceived the project. M.G.E and S.T.K. created the gene sequences. M.G.E. and A.T.P. engineered proteins and characterized their properties. M.G.E. and M.M.M. performed flow cytometry and *in vivo* binding assays. All authors analyzed data. M.G.E. and R.A.C. wrote the manuscript. H.G.G. edited the manuscript.

## Methods

### Chemicals and enzymes

All reagents used were of the highest available purity. Synthetic oligonucleotides were purchased from Eurofins MWG Operon. Restriction enzymes and DNA-modifying enzymes were purchased from New England Biolabs. All aqueous solutions were prepared using water purified with a Barnstead Nanopure Diamond system.

### Mutagenesis and cloning

Codon-optimized (*E. coli*) and his-tagged (N-terminus) sequences for gSTEP0 and STEPtag (Supplementary Table 1) were purchased from ATUM. Truncation mutants of gSTEP0 (T1–T4) were obtained by polymerase chain reaction amplification of the appropriate region of the gene, while mutants with added linkers (L1–L5) or alternate Bim peptides (hBim, XXA1, XXA4, G2gE, Y4eK) were generated using splicing by overlap extension (SOE) mutagenesis.^*36*^ The combinatorial linker saturation library was generated by SOE mutagenesis of gSTEP0-T1-L4 using oligonucleotides containing four consecutive NNS degenerate codons, one for every position of the linker sequence. All sequences were subcloned into pET11a vectors (Novagen) *via* the *Nde*I/*Bam*HI restriction sites. Gene constructs for live-cell experiments (i.e., flow cytometry and *in vivo* binding assays) were subcloned via *Nco*I/*Eco*RI or *Hind*III/*Bam*HI into either the pBAD/His A (Invitrogen) or pZA23MCS (EXPRESSYS) vectors for inducible or constitutive expression, respectively. *Aequorea victoria* EGFP [Genbank AAB02572] was cloned into pBAD/His A using *Xho*I/*Eco*RI, which added the pBAD His tag/Xpress^™^ Epitope/EK site to the N-terminus. His-tagged (C-terminus) *Thermoascus aurantiacus* xylanase 10A (TAX, UniProtKB: P23360) in which the two catalytic residues were mutated to alanine (E157A/E263A) cloned into a pET11a vector via *Nde*I/*Bam*HI was a gift from Stephen L. Mayo.^*37*^ TAX-L10-STEPtag and STEPtag-L10-TAX constructs were generated using SOE mutagenesis and cloned into pET11a vectors as described above. His-tagged (N-terminus) sfGFP cloned into a pBAD vector (pBAD-sfGFP)^*32*^ was a gift from Michael Davidson & Geoffrey Waldo (Addgene plasmid #54519; http://n2t.net/addgene:54519; RRID: Addgene_54519). All constructs were verified by sequencing the entire open reading frame (see Supplementary Table 1 for amino-acid sequences), and transformed into either BL21-Gold(DE3) (Agilent) or TOP10 (Thermo Fisher) chemically-competent *E. coli* cells for pET11a, or pBAD and pZA vectors, respectively.

### Protein expression and purification

Transformed *E. coli* cells harboring expression vectors were grown in 500 mL lysogeny broth (LB) supplemented with 100 µg mL^−1^ ampicillin at 37°C with shaking. When an OD600 of 0.6–0.8 was reached, protein expression was induced by addition of 1 mM isopropyl β-D-1-thiogalactopyranoside (pET11a vectors) or 0.2% arabinose (pBAD vectors). Following overnight incubation at 16°C with shaking, cells were harvested by centrifugation and lysed with an EmulsiFlex-B15 cell disruptor (Avestin). Following removal of cellular debris by centrifugation, proteins were extracted and purified by immobilized metal affinity chromatography using Profinity IMAC resin (Bio-Rad) in a gravity flow column according to the manufacturer’s protocol. Eluted proteins were exchanged into 20 mM sodium phosphate buffer containing 50 mM NaCl (pH 7.4) and concentrated using Amicon Ultra-15 centrifugal filters with a molecular weight cut-off of 3 kDa (Millipore) for STEPtag, or Microsep Advance centrifugal filters with a molecular weight cut-off of 10 kDa (Pall) for all other proteins. Purified proteins were quantified by measuring absorbance at 280 nm in a 1-cm quartz cuvette with a SpectraMax Plus384 microplate spectrophotometer (Molecular Devices), and applying Beer-Lambert’s law using extinction coefficients calculated with the ProtParam tool (https://web.expasy.org/protparam/).

### In vitro binding assays

All fluor escence measurements were performed in triplicate wells of Fluotrac 96-well plates (Greiner Bio-One) on a Tecan Infinite M1000 plate reader using 75 nM of each gSTEP variant in 20 mM sodium phosphate buffer containing 50 mM NaCl (pH 7.4). To calculate *K*_d_ and ΔF/F_0_ values, gSTEP fluorescence intensity (λ_ex_ = 485 nm, λ_em_ = 515 nm) as a function of STEPtag, TAX-L10-STEPtag, STEPtag-L10-TAX, or control protein concentration (e.g., bovine serum albumin [Bio-Rad] or an inactive mutant of *Thermoascus aurantiacus* xylanase 10A purified as described above^*37*^) was fit to the Hill equation, accounting for ligand depletion^*38*^:

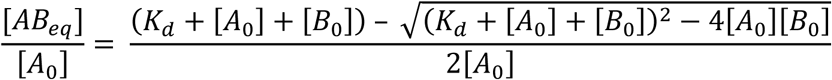

where *A* (gSTEP variants) and *B* (STEPtag, TAX-L10-STEPtag, or STEPtag-L10-TAX) are the two binding proteins, and [*A*_0_] and [*B*_0_] are the initial concentrations of each protein. [*AB*_eq_] is the equilibrium concentration of the bound complex.

### Fluorescence-activated cell sorting

To improve the signal-to-noise ratio in live cells, we aimed to isolate gSTEP0-T1-L4 variants that gave the brightest fluorescence from the linker saturation library. To do so, we transformed the gSTEP0-T1-L4 mutant library into E. cloni^®^ Elite electrocompetent *E. coli* cells (Lucigen), which were plated on LB agar supplemented with 100 µg mL^−1^ ampicillin. Following overnight incubation at 37°C, a total of 10^5^ colonies from multiple agar plates were collected, pooled together, and cultured overnight in 10 mL LB supplemented with ampicillin. Following extraction of plasmid DNA from this culture, the library was transformed into BL21-Gold(DE3) electrocompetent *E. coli* cells, and plated on LB agar supplemented with ampicillin. From these plates, 10^5^ colonies were collected, pooled together, and cultured overnight in 10 mL LB supplemented with ampicillin. This bacterial culture was diluted 100-fold into fresh LB supplemented with ampicillin and grown to an OD600 of 0.5–0.9. Because the leaky expression of the T7 RNA polymerase in BL21-Gold(DE3) provided sufficient quantities of protein to screen, the cells were not further induced with isopropyl β-D-1-thiogalactopyranoside to limit their metabolic burden. After growth, cells were centrifuged and pellets were washed twice with filter-sterilized 20 mM sodium phosphate buffer containing 50 mM NaCl (pH 7.4). Resuspended cells were diluted in this buffer to a concentration of approximately 5 × 10^7^ colony forming units per mL.^*39*^ The cells were then filtered twice using a 40-µm Falcon Cell Strainer (Fisher) to remove large particulates. Fluorescence-activated cell sorting was performed on a MoFlo AstriosEQ Cell Sorter (Beckman Coulter) using a 488 nm laser for excitation and a 513/26 nm filter for detecting fluorescence emission. Data analysis was performed with the FlowJo software package (BD). This process was repeated twice in succession, collecting 20000 of the brightest cells each time.

The collected cells were used to inoculate 50 mL of fresh LB supplemented with ampicillin, and grown overnight at 37°C with shaking. This culture was used to streak an LB agar plate supplemented with ampicillin. From this plate, 96 colonies were picked into individual wells of a Nunc V96 MicroWell polypropylene plate containing 200 µL of LB with 100 µg mL^−1^ ampicillin supplemented with 10% glycerol. The plate was covered with a sterile gas permeable rayon film (VWR) and incubated overnight at 37°C with shaking. After incubation, the mother plate was used to inoculate duplicate Nunc V96 MicroWell polypropylene plates (daughter plates) containing 250 µL of LB with 100 µg mL^−1^ ampicillin per well. Daughter plates were sealed with rayon film and incubated overnight (37°C, 250 rpm shaking). After incubation, the cells were harvested by centrifugation and the pellets were washed twice with phosphate buffered saline. These pellets were resuspended and lysed in 100 µL of Bugbuster protein extraction reagent (Millipore) containing 5 U mL^−1^ Benzonase nuclease (Millipore) and 1 mg ml^−1^ hen egg white lysozyme (Omnipure). Following centrifugation to remove cellular debris, the clarified lysate (30 µL) was transferred to a Fluotrac 96-well plate (Greiner Bio-One) for screening. To each 30-µL lysate containing a different gSTEP0-T1-L4 variant, 150 µL of 20 mM sodium phosphate buffer containing 50 mM NaCl (pH 7.4) and 0 or 9 µM purified STEPtag was added. Fluorescence was measured with a Tecan Infinite M1000 plate reader. Emission spectra (λ_ex_ = 485 nm) were measured from 500 nm to 560 nm. From these spectra, ΔF/F_0_ was calculated for each protein variant, and the one with the best response (gSTEP1) was analyzed further.

### Rapid-mixing stopped-flow kinetics

Measurements were performed using an RSM 1000 UV/Vis rapid-scanning spectrophotometer (Olis) equipped with a 1.24-mm-slit fixed disk for single wavelength measurements, and plane gratings with 400 lines mm^−1^ and a 500 nm blaze wavelength. All other fixed slits were set to 3.16 mm to maximize signal. Purified gSTEP1 (1 µM) and STEPtag (5 µM) were loaded into the spectrophotometer, which was kept at 37°C using a temperature control unit (Julabo). 300 µL of each sample was pumped into the mixing chamber, and the fluorescence was measured (λ_ex_ = 485 nm, λ_em_ = 515 nm). For each combination of samples, the dead volume was cleared prior to data collection. Control experiments were performed to confirm that fluorescence increase was due to binding of gSTEP1 to STEPtag (Supplementary Figure 5). The data was fit to the integrated rate equation, accounting for ligand depletion^*38*^,

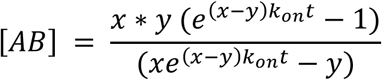

where *A* and *B* are the two binding proteins (gSTEP1 and STEPtag), *x* = [*AB*_eq_], *y* = [*A*_0_] [*B*_0_] / [AB_eq_], and *t* is the time.

### Flow cytometry

TOP10 *E. coli* cells (Invitrogen) transformed with pZA-gSTEP1 and/or pBAD-STEPtag vectors were cultured in 50 mL LB supplemented with 100 µg mL^−1^ ampicillin (for cells containing pBAD) and/or 50 µg mL^−1^ kanamycin (for cells containing pZA). Cells were grown with shaking at 37°C to an OD600 of 0.4–0.8, then the culture containing both pBAD-STEPtag and pZA-gSTEP1 was split equally into two flasks, one to be induced and the other to be left uninduced. Following induction of cells containing pBAD vectors with 0.2% arabinose, cultures were incubated for an additional 60 minutes at 37°C with shaking. Cells were then harvested by centrifugation, and prepared for flow cytometry as described in the cell sorting protocol above. Two biological replicates of flow cytometry measurements were performed using a Gallios flow cytometer (Beckman Coulter), set to detect either 10000 or 100000 events per run. Fluorescence was detected with a 525/40 filter (λ_ex_ = 488 nm), and data analysis was performed using the Kaluza software package (Beckman Coulter).

### In vivo binding assays

TOP10 *E. coli* cells transformed with the appropriate vectors were cultured as described for the flow cytometry experiments above. Cells were grown with shaking at 37°C to an OD600 of 0.6–1.1, after which 200 µL of each culture was transferred to a Fluotrac 96-well plate (Greiner Bio-One) in triplicate wells. Fluorescence measurements were recorded on an Infinite M1000 microplate reader equipped with an injector module (Tecan), preheated to 37°C (λ_ex_ = 488 nm, λ_em_ = 514 nm). Measurements were taken every 2 minutes for 10 minutes, shaking the plate before each measurement, then protein expression was induced by injecting 12 µL of 8% arabinose into the wells (final concentration of 0.45%), followed by 3 seconds of shaking and 2 seconds of settle time. Fluorescence was measured every 2–6 seconds for an additional 20 or 40 minutes (Supplementary Figure 4).

## Abbreviations

GFP: green fluorescent protein
STEP: sensor for transiently-expressed proteins
cpGFP: circularly permuted green fluorescent protein
gSTEP: green fluorescent sensor for transiently expressed proteins
EGFP: enhanced green fluorescent protein
sfGFP: superfolder green fluorescent protein
SOE: splicing by overlap extension
LB: lysogeny broth.

## Graphical abstract

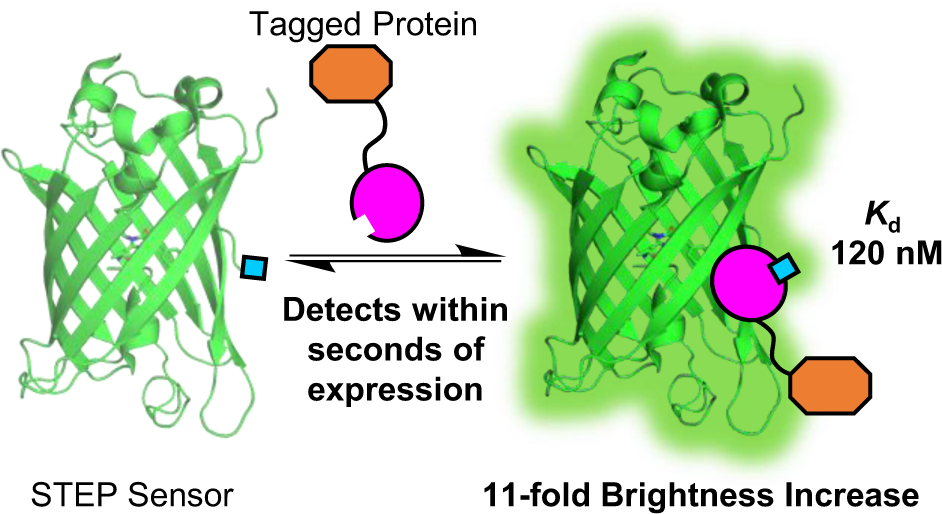

