## Supplementary Materials for "Genetically-encoded fluorescent biosensor for rapid detection of protein expression"

**This file includes:**

Supplementary Tables 1–2
Supplementary Figures 1–5

**Supplementary Table 1.** Amino-acid sequences of STEP biosensor and fluorescent protein constructs.

| **Protein** | **Molecular Weight (kDa)** | **Sequence**[a] |
|---|---|---|
| **gSTEP1** | 31.8 | MHHHHHHDLRPEIWIAQELRRIGDEFNAYYARRHKIRLENVYIKADKQKN<br>GIKANFKIRHNIEDGGVQLAYHYQQNTPIGDGPVLLPDNHYLSVQSKLSK<br>DPNEKRDHMVLLEFVTAAGITLGMDELYKGGTGGSMVSKGEELFTGVVPI<br>LVELDGDVNGHKFSVSGEGEGDATYGKLTLKFICTTGKLPVPWPTLVTTL<br>TYGVQCFSRYPDHMKQHDFFKSAMPEGYIQERTIFFKDDGNYKTRAEVKF<br>EGDTLVNRIELKGIDFKEDGNILGHKLEY |
| **STEPtag** | 15.5 | MHHHHHHREVIPMAAVKQALREAGDEFELRYRRAFSDLTSQLHITPGTAY<br>QSFEQVVNELFRDGVNWGRIVAFFSFGGALCVESVDKEMQVLVSRIAAWM<br>ATYLNDHLEPWIQENGGWDTFVELYGNNAAAESRK |
| **cpGFP** | 29.0 | MHHHHHHENLYFQGLENVYIKADKQKNGIKANFKIRHNIEDGGVQLAYHY<br>QQNTPIGDGPVLLPDNHYLSVQSKLSKDPNEKRDHMVLLEFVTAAGITLG<br>MDELYKGGTGGSMVSKGEELFTGVVPILVELDGDVNGHKFSVSGEGEGDA<br>TYGKLTLKFICTTGKLPVPWPTLVTTLTYGVQCFSRYPDHMKQHDFFKSA<br>MPEGYIQERTIFFKDDGNYKTRAEVKFEGDTLVNRIELKGIDFKEDGNIL<br>GHKLEYN |
| **gSTEP0** | 31.6 | MHHHHHHDLRPEIRIAQELRRIGDEFNETYTRRLENVYIKADKQKNGIKA<br>NFKIRHNIEDGGVQLAYHYQQNTPIGDGPVLLPDNHYLSVQSKLSKDPNE<br>KRDHMVLLEFVTAAGITLGMDELYKGGTGGSMVSKGEELFTGVVPILVEL<br>DGDVNGHKFSVSGEGEGDATYGKLTLKFICTTGKLPVPWPTLVTTLTYGV<br>QCFSRYPDHMKQHDFFKSAMPEGYIQERTIFFKDDGNYKTRAEVKFEGDT<br>LVNRIELKGIDFKEDGNILGHKLEYNTR |
| **gSTEP0-T1** | 31.2 | MHHHHHHDLRPEIRIAQELRRIGDEFNETYTRRLENVYIKADKQKNGIKA<br>NFKIRHNIEDGGVQLAYHYQQNTPIGDGPVLLPDNHYLSVQSKLSKDPNE<br>KRDHMVLLEFVTAAGITLGMDELYKGGTGGSMVSKGEELFTGVVPILVEL<br>DGDVNGHKFSVSGEGEGDATYGKLTLKFICTTGKLPVPWPTLVTTLTYGV<br>QCFSRYPDHMKQHDFFKSAMPEGYIQERTIFFKDDGNYKTRAEVKFEGDT<br>LVNRIELKGIDFKEDGNILGHKLEY |
| **gSTEP0-T2** | 31.1 | MHHHHHHDLRPEIRIAQELRRIGDEFNETYTRRLENVYIKADKQKNGIKA<br>NFKIRHNIEDGGVQLAYHYQQNTPIGDGPVLLPDNHYLSVQSKLSKDPNE<br>KRDHMVLLEFVTAAGITLGMDELYKGGTGGSMVSKGEELFTGVVPILVEL<br>DGDVNGHKFSVSGEGEGDATYGKLTLKFICTTGKLPVPWPTLVTTLTYGV<br>QCFSRYPDHMKQHDFFKSAMPEGYIQERTIFFKDDGNYKTRAEVKFEGDT<br>LVNRIELKGIDFKEDGNILGHKLE |
| **gSTEP0-T3** | 30.9 | MHHHHHHDLRPEIRIAQELRRIGDEFNETYTRRLENVYIKADKQKNGIKA<br>NFKIRHNIEDGGVQLAYHYQQNTPIGDGPVLLPDNHYLSVQSKLSKDPNE<br>KRDHMVLLEFVTAAGITLGMDELYKGGTGGSMVSKGEELFTGVVPILVEL<br>DGDVNGHKFSVSGEGEGDATYGKLTLKFICTTGKLPVPWPTLVTTLTYGV<br>QCFSRYPDHMKQHDFFKSAMPEGYIQERTIFFKDDGNYKTRAEVKFEGDT<br>LVNRIELKGIDFKEDGNILGHKL |
| **gSTEP0-T4** | 30.8 | MHHHHHHDLRPEIRIAQELRRIGDEFNETYTRRLENVYIKADKQKNGIKA<br>NFKIRHNIEDGGVQLAYHYQQNTPIGDGPVLLPDNHYLSVQSKLSKDPNE<br>KRDHMVLLEFVTAAGITLGMDELYKGGTGGSMVSKGEELFTGVVPILVEL<br>DGDVNGHKFSVSGEGEGDATYGKLTLKFICTTGKLPVPWPTLVTTLTYGV<br>QCFSRYPDHMKQHDFFKSAMPEGYIQERTIFFKDDGNYKTRAEVKFEGDT<br>LVNRIELKGIDFKEDGNILGHK |
| **gSTEP0-T1-hBim** | 31.2 | MHHHHHHDLRPEIWIAQELRRIGDEFNAYYARRLENVYIKADKQKNGIKA<br>NFKIRHNIEDGGVQLAYHYQQNTPIGDGPVLLPDNHYLSVQSKLSKDPNE<br>KRDHMVLLEFVTAAGITLGMDELYKGGTGGSMVSKGEELFTGVVPILVEL<br>DGDVNGHKFSVSGEGEGDATYGKLTLKFICTTGKLPVPWPTLVTTLTYGV<br>QCFSRYPDHMKQHDFFKSAMPEGYIQERTIFFKDDGNYKTRAEVKFEGDT<br>LVNRIELKGIDFKEDGNILGHKLEY |

| | | |
|---|---|---|
| **gSTEP0-T1-XXA1[b]** | 31.2 | MHHHHHHDLRPEIWYAQGLKRFGDEFNAYYARRLENVYIKADKQKNGIKANFKIRHNIEDGGVQLAYHYQQNTPIGDGPVLLPDNHYLSVQSKLSKDPNEKRDHMVLLEFVTAAGITLGMDELYKGGTGGSMVSKGEELFTGVVPILVELDGDVNGHKFSVSGEGEGDATYGKLTLKFICTTGKLPVPWPTLVTTLTYGVQCFSRYPDHMKQHDFFKSAMPEGYIQERTIFFKDDGNYKTRAEVKFEGDTLVNRIELKGIDFKEDGNILGHKLEY |
| **gSTEP0-T1-XXA4[b]** | 31.3 | MHHHHHHDLRPEIWYAQWLKRFGDQFNAYYARRLENVYIKADKQKNGIKANFKIRHNIEDGGVQLAYHYQQNTPIGDGPVLLPDNHYLSVQSKLSKDPNEKRDHMVLLEFVTAAGITLGMDELYKGGTGGSMVSKGEELFTGVVPILVELDGDVNGHKFSVSGEGEGDATYGKLTLKFICTTGKLPVPWPTLVTTLTYGVQCFSRYPDHMKQHDFFKSAMPEGYIQERTIFFKDDGNYKTRAEVKFEGDTLVNRIELKGIDFKEDGNILGHKLEY |
| **gSTEP0-T1-G2gE[b]** | 31.3 | MHHHHHHDLRPEIWYAQELKRFGDEFNAYYARRLENVYIKADKQKNGIKANFKIRHNIEDGGVQLAYHYQQNTPIGDGPVLLPDNHYLSVQSKLSKDPNEKRDHMVLLEFVTAAGITLGMDELYKGGTGGSMVSKGEELFTGVVPILVELDGDVNGHKFSVSGEGEGDATYGKLTLKFICTTGKLPVPWPTLVTTLTYGVQCFSRYPDHMKQHDFFKSAMPEGYIQERTIFFKDDGNYKTRAEVKFEGDTLVNRIELKGIDFKEDGNILGHKLEY |
| **gSTEP0-T1-Y4eK[b]** | 31.2 | MHHHHHHDLRPEIWYAQGLKRFGDEFNAYKARRLENVYIKADKQKNGIKANFKIRHNIEDGGVQLAYHYQQNTPIGDGPVLLPDNHYLSVQSKLSKDPNEKRDHMVLLEFVTAAGITLGMDELYKGGTGGSMVSKGEELFTGVVPILVELDGDVNGHKFSVSGEGEGDATYGKLTLKFICTTGKLPVPWPTLVTTLTYGVQCFSRYPDHMKQHDFFKSAMPEGYIQERTIFFKDDGNYKTRAEVKFEGDTLVNRIELKGIDFKEDGNILGHKLEY |
| **gSTEP0-T1-L1** | 31.3 | MHHHHHHDLRPEIRIAQELRRIGDEFNETYTRRGLENVYIKADKQKNGIKANFKIRHNIEDGGVQLAYHYQQNTPIGDGPVLLPDNHYLSVQSKLSKDPNEKRDHMVLLEFVTAAGITLGMDELYKGGTGGSMVSKGEELFTGVVPILVELDGDVNGHKFSVSGEGEGDATYGKLTLKFICTTGKLPVPWPTLVTTLTYGVQCFSRYPDHMKQHDFFKSAMPEGYIQERTIFFKDDGNYKTRAEVKFEGDTLVNRIELKGIDFKEDGNILGHKLEY |
| **gSTEP0-T1-L2** | 31.4 | MHHHHHHDLRPEIRIAQELRRIGDEFNETYTRRGSLENVYIKADKQKNGIKANFKIRHNIEDGGVQLAYHYQQNTPIGDGPVLLPDNHYLSVQSKLSKDPNEKRDHMVLLEFVTAAGITLGMDELYKGGTGGSMVSKGEELFTGVVPILVELDGDVNGHKFSVSGEGEGDATYGKLTLKFICTTGKLPVPWPTLVTTLTYGVQCFSRYPDHMKQHDFFKSAMPEGYIQERTIFFKDDGNYKTRAEVKFEGDTLVNRIELKGIDFKEDGNILGHKLEY |
| **gSTEP0-T1-L3** | 31.4 | MHHHHHHDLRPEIRIAQELRRIGDEFNETYTRRGSGLENVYIKADKQKNGIKANFKIRHNIEDGGVQLAYHYQQNTPIGDGPVLLPDNHYLSVQSKLSKDPNEKRDHMVLLEFVTAAGITLGMDELYKGGTGGSMVSKGEELFTGVVPILVELDGDVNGHKFSVSGEGEGDATYGKLTLKFICTTGKLPVPWPTLVTTLTYGVQCFSRYPDHMKQHDFFKSAMPEGYIQERTIFFKDDGNYKTRAEVKFEGDTLVNRIELKGIDFKEDGNILGHKLEY |
| **gSTEP0-T1-L4** | 31.5 | MHHHHHHDLRPEIRIAQELRRIGDEFNETYTRRGSGSLENVYIKADKQKNGIKANFKIRHNIEDGGVQLAYHYQQNTPIGDGPVLLPDNHYLSVQSKLSKDPNEKRDHMVLLEFVTAAGITLGMDELYKGGTGGSMVSKGEELFTGVVPILVELDGDVNGHKFSVSGEGEGDATYGKLTLKFICTTGKLPVPWPTLVTTLTYGVQCFSRYPDHMKQHDFFKSAMPEGYIQERTIFFKDDGNYKTRAEVKFEGDTLVNRIELKGIDFKEDGNILGHKLEY |
| **gSTEP0-T1-L4-hBim** | 31.5 | MHHHHHHDLRPEIWIAQELRRIGDEFNAYYARRGSGSLENVYIKADKQKNGIKANFKIRHNIEDGGVQLAYHYQQNTPIGDGPVLLPDNHYLSVQSKLSKDPNEKRDHMVLLEFVTAAGITLGMDELYKGGTGGSMVSKGEELFTGVVPILVELDGDVNGHKFSVSGEGEGDATYGKLTLKFICTTGKLPVPWPTLVTTLTYGVQCFSRYPDHMKQHDFFKSAMPEGYIQERTIFFKDDGNYKTRAEVKFEGDTLVNRIELKGIDFKEDGNILGHKLEY |

| | | |
|---|---|---|
| **gSTEP0-T1-L5** | 31.6 | MHHHHHHDLRPEIRIAQELRRIGDEFNETYTRRGSGSGLENVYIKADKQK<br>NGIKANFKIRHNIEDGGVQLAYHYQQNTPIGDGPVLLPDNHYLSVQSKLS<br>KDPNEKRDHMVLLEFVTAAGITLGMDELYKGGTGGSMVSKGEELFTGVVP<br>ILVELDGDVNGHKFSVSGEGEGDATYGKLTLKFICTTGKLPVPWPTLVTT<br>LTYGVQCFSRYPDHMKQHDFFKSAMPEGYIQERTIFFKDDGNYKTRAEVK<br>FEGDTLVNRIELKGIDFKEDGNILGHKLEY |
| **STEPtag-L10-TAX** | 49.8 | MHHHHHHREVIPMAAVKQALREAGDEFELRYRRAFSDLTSQLHITPGTAY<br>QSFEQVVNELFRDGVNWGRIVAFFSFGGALCVESVDKEMQVLVSRIAAWM<br>ATYLNDHLEPWIQENGGWDTFVELYGNNAAAESRKGSSGGGGSGGMAEAA<br>QSVDQLIKARGKVYFGVATDQNRLTTGKNAAIIQADFGQVTPENSMKWDA<br>TEPSQGNFNFAGADYLVNWAQQNGKLIRGHTLVWHSQLPSWVSSITDKNT<br>LTNVMKNHITTLMTRYKGKIRAWDVVNAAFNEDGSLRQTVFLNVIGEDYI<br>PIAFQTARAADPNAKLYINDYNLDSASYPKTQAIVNRVKQWRAAGVPIDG<br>IGSQTHLSAGQGAGVLQALPLLASAGTPEVAITALDVAGASPTDYVNVVN<br>ACLNVQSCVGITVWGVADPDSWRASTTPLLFDGNFNPKPAYNAIVQDLQQ<br>GSIEGRG |
| **TAX-L10-STEPtag** | 49.6 | MAEAAQSVDQLIKARGKVYFGVATDQNRLTTGKNAAIIQADFGQVTPENS<br>MKWDATEPSQGNFNFAGADYLVNWAQQNGKLIRGHTLVWHSQLPSWVSSI<br>TDKNTLTNVMKNHITTLMTRYKGKIRAWDVVNAAFNEDGSLRQTVFLNVI<br>GEDYIPIAFQTARAADPNAKLYINDYNLDSASYPKTQAIVNRVKQWRAAG<br>VPIDGIGSQTHLSAGQGAGVLQALPLLASAGTPEVAITALDVAGASPTDY<br>VNVVNACLNVQSCVGITVWGVADPDSWRASTTPLLFDGNFNPKPAYNAIV<br>QDLQQGSIEGRGGSSGGGGSGGREVIPMAAVKQALREAGDEFELRYRRAF<br>SDLTSQLHITPGTAYQSFEQVVNELFRDGVNWGRIVAFFSFGGALCVESV<br>DKEMQVLVSRIAAWMATYLNDHLEPWIQENGGWDTFVELYGNNAAAESRK<br>HHHHHH |
| **EGFP**[c] | 31.1 | MGGSHHHHHHGMASMTGGQQMGRDLYDDDDKDRWGSELEGSKGEELFTGV<br>VPILVELDGDVNGHKFSVSGEGEGDATYGKLTLKFICTTGKLPVPWPTLV<br>TTLTYGVQCFSRYPDHMKQHDFFKSAMPEGYVQERTIFFKDDGNYKTRAE<br>VKFEGDTLVNRIELKGIDFKEDGNILGHKLEYNYNSHNVYIMADKQKNGI<br>KVNFKIRHNIEDGSVQLADHYQQNTPIGDGPVLLPDNHYLSTQSALSKDP<br>NEKRDHMVLLEFVTAAGITLGMDELYK |

[a] Sequences of fluorescent proteins, Bim peptides, Bcl-$x_L$, *Thermoascus aurantiacus* xylanase 10A (TAX), and linkers are highlighted in green, cyan, magenta, orange, and grey, respectively.
[b] Sequences contain engineered variants of the human Bim peptide (from Dutta *et al.*, *Journal of Molecular Biology* 2015, 427, 1241–1253). gSTEP0-T1-Y4eK contains the XXA1_Y4eK variant.
[c] The EGFP sequence used here includes a 40-amino-acid, N-terminal tag from the pBAD/HisA vector.

**Supplementary Table 2. Engineering of gSTEP1**

| Sensor | $\lambda_{ex}$ (nm) | $\lambda_{em}$ (nm) | $K_d$ (nM) | $\Delta F/F_0$ |
|---|---|---|---|---|
| Initial prototype | | | | |
| gSTEP0[a] | 496 | 513 | 250 ± 40 | 1.4 ± 0.1 |
| Truncated variants | | | | |
| gSTEP0-T1[a] | 504 | 515 | 210 ± 80 | 2.1 ± 0.4 |
| gSTEP0-T2 | N.D. | N.D. | 200 | 1.8 |
| gSTEP0-T3 | N.D. | N.D. | 230 | 1.9 |
| gSTEP0-T4[b] | N.D. | N.D. | N.D. | N.D. |
| Bim peptide variants | | | | |
| gSTEP0-T1-hBim[a] | 502 | 514 | 170 ± 40 | 3.3 ± 0.6 |
| gSTEP0-T1-XXA1 | 502 | 515 | 190 | 2.3 |
| gSTEP0-T1-XXA4[a] | 500 | 513 | 280 ± 50 | 1.3 ± 0.1 |
| gSTEP0-T1-G2gE[c] | 503 | 513 | 190 ± 20 | 2.4 ± 0.2 |
| gSTEP0-T1-Y4eK[a] | 504 | 515 | 260 ± 20 | 1.4 ± 0.1 |
| Linker variants | | | | |
| gSTEP0-T1-L1[d] | 500 | 514 | N.D. | N.D. |
| gSTEP0-T1-L2[c] | 499 | 514 | 400 ± 100 | 1.3 ± 0.1 |
| gSTEP0-T1-L3 | 501 | 513 | 170 | 1.4 |
| gSTEP0-T1-L4[a] | 502 | 515 | 140 ± 20 | 2.4 ± 0.2 |
| gSTEP0-T1-L4-hBim[c] | 500 | 515 | 290 ± 80 | 1.0 ± 0.1 |
| gSTEP0-T1-L5 | 498 | 513 | 500 | 1.0 |
| Final variant | | | | |
| gSTEP1[f] | 504 | 515 | 120 ± 20 | 3.4 ± 0.4 |

N.D. indicates not determined.
[a] n = 3 biological replicates, fit value ± 95% confidence interval
[b] No binding curves were performed for gSTEP0-T4 due to low fluorescence
[c] n = 2 biological replicates, fit value ± 95% confidence interval
[d] No change in the fluorescence of gSTEP0-T1-L1 was observed upon addition of 10 µM STEPtag
[e] n = 5 biological replicates, fit value ± 95% confidence interval
[f] n = 6 biological replicates, fit value ± 95% confidence interval

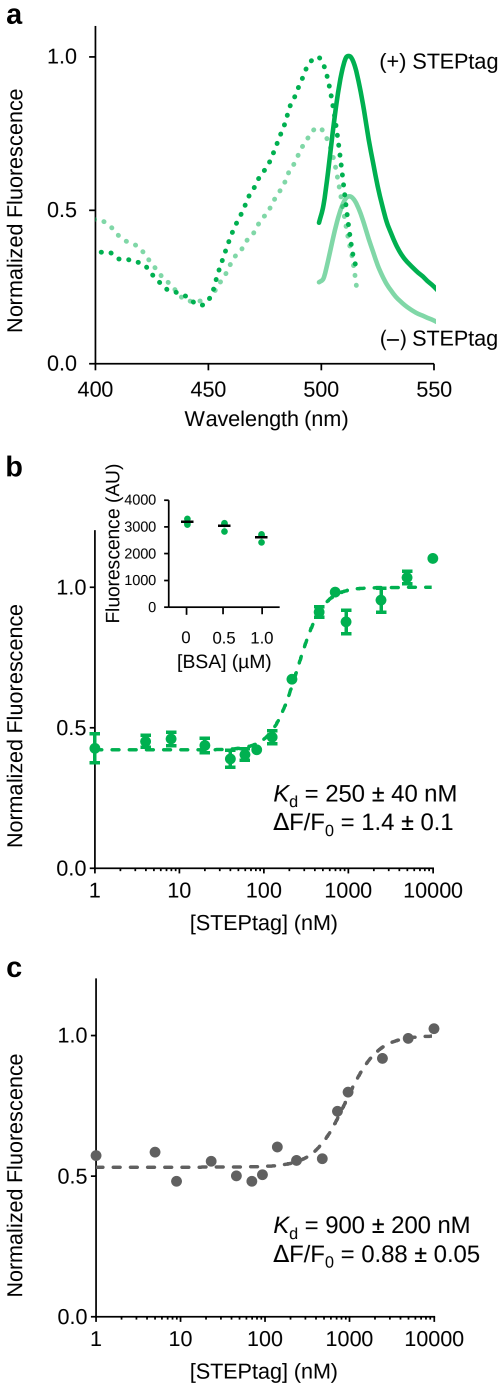


**Supplementary Figure 1.** ***In vitro* characterization of gSTEP0. a,** Excitation ($\lambda_{em}$ = 550 nm) and emission ($\lambda_{ex}$ = 485 nm) spectra of gSTEP0 (75 nM) alone or in the presence of saturating STEPtag (10 µM) are shown as dotted or full lines, respectively. **b,** Binding curve of gSTEP0 with STEPtag. The dashed line represents a fit of the Hill equation to the data (Hill coefficient = 2.9). Data points represent mean ± SEM of two biological replicates, each performed in triplicate. Fluorescence values are normalized to the maximum intensity fit. Inset shows the fluorescence intensity at 550 nm ($\lambda_{ex}$ = 495 nm) of 75 nM gSTEP0 in the presence or absence of bovine serum albumin (BSA). N = 3 technical replicates, mean shown as black bar. **c,** Binding curve of cpGFP with STEPtag. The dashed line represents a fit of the Hill equation to the data (Hill coefficient = 2.3). Data points represent mean of triplicate measurements. Fluorescence values were normalized to the maximum intensity fit.

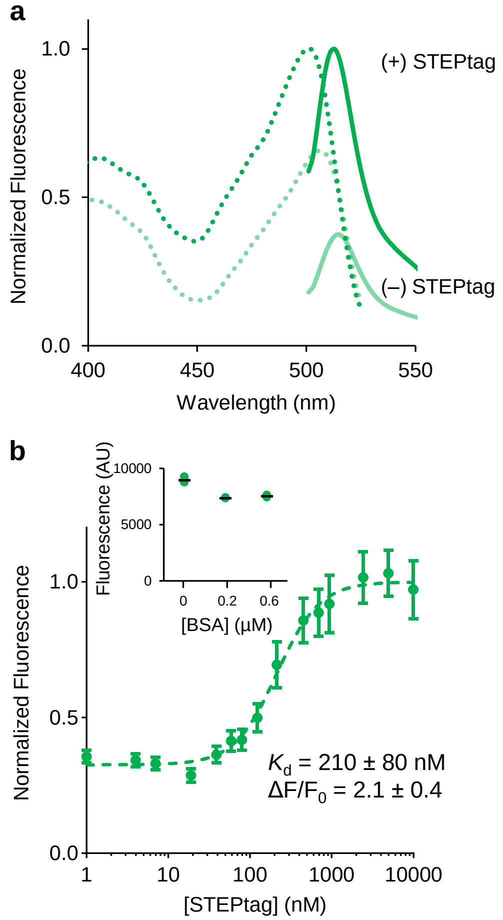


**Supplementary Figure 2. *In vitro* characterization of gSTEP0-T1. a,** Excitation ($\lambda_{em}$ = 550 nm) and emission ($\lambda_{ex}$ = 485 nm) spectra of gSTEP0-T1 (75 nM) alone or in the presence of saturating STEPtag (10 µM) are shown as dotted or full lines, respectively. **b,** Binding curve of gSTEP0-T1 with STEPtag. The dashed line represents a fit of the Hill equation to the data (Hill coefficient = 1.7). Data points represent mean ± SEM of three biological replicates, each performed in triplicate. Fluorescence values were normalized to the maximum intensity fit. Inset shows the fluorescence intensity at 530 nm ($\lambda_{ex}$ = 488 nm) of three technical replicates of 75 nM gSTEP0-T1, in the presence or absence of bovine serum albumin (BSA). The mean value is indicated by a black bar.

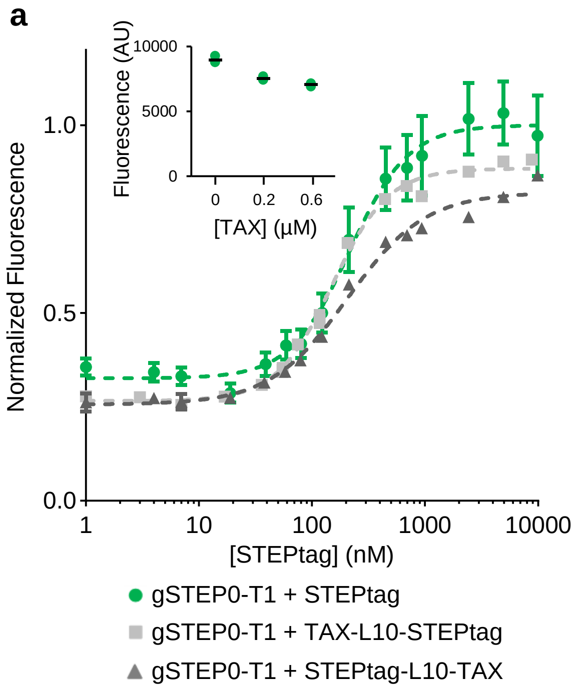


**b**

| **Binding Partner** | ***K*$_d$ (nM)** | **ΔF/F$_0$** |
|---|---|---|
| STEPtag [a] | 210 ± 80 | 2.1 ± 0.4 |
| TAX-L10-STEPtag [b] | 150 ± 10 | 2.3 ± 0.1 |
| STEPtag-L10-TAX [c] | 200 ± 30 | 2.2 ± 0.2 |

[a] Data obtained from three biological replicates
[b] Data obtained from one biological replicate
[c] Data obtained from two biological replicates
In all cases, error shown is the 95% confidence interval

**Supplementary Figure 3. Binding affinity and fluorescence response of gSTEP0-T1 is not substantially affected by fusion of STEPtag to a protein of interest. a**, Binding curves of gSTEP0-T1 with STEPtag alone or fused at the N- or C-terminus of the TAX protein via a 10 amino-acid linker (L10). TAX is the xylanase 10A enzyme from *Thermoascus aurantiacus*, a 36 kDa globular protein. Dashed lines represent fits of the Hill equation to the data (Hill coefficients of 1.7, 1.8, and 1.3 for STEPtag, TAX-L10-STEPtag, STEPtag-L10-TAX, respectively). For STEPtag, n = 3 biological replicates, mean ± SEM. For TAX-L10-STEPtag, n = 1 biological replicate. For STEPtag-L10-TAX, n = 2 biological replicates, mean ± SEM. Inset shows the fluorescence intensity at 530 nm ($\lambda_{ex}$ = 488 nm) of three technical replicates of 75 nM gSTEP0-T1, in the presence or absence of TAX (mean represented by a black bar). As shown, untagged TAX does not cause an increase of gSTEP0-T1 fluorescence. **b,** Binding affinity ($K_d$) and fluorescence response ($\Delta F/F_0$) of 75 nM gSTEP0-T1 with various binding partners. With a 10 amino-acid linker (L10), binding of STEPtag to gSTEP0-T1 is not significantly affected by the presence of the TAX protein.

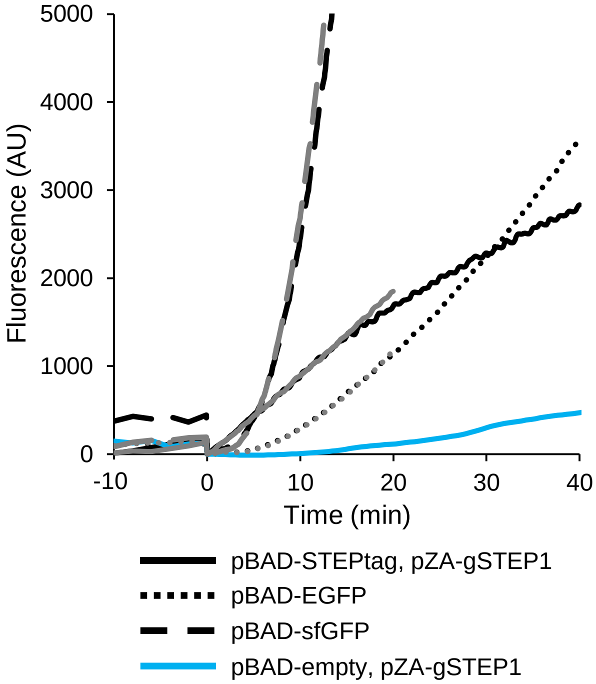


**Supplementary Figure 4. *In vivo* binding assay.** Time course of protein expression in live *E. coli*. Cells were grown to the end of the exponential growth phase (OD600 = 1.1), then fluorescence was measured every two minutes for 10 minutes, at which point expression of STEPtag (for cells constitutively expressing gSTEP1), EGFP, or sfGFP was induced with 0.45% arabinose (t = 0 min). Fluorescence was then measured as rapidly as possible. Two biological replicates are shown (black, stopped after 40 min, and grey, stopped after 20 min), and one control experiment with cells containing pZA-gSTEP1 and an empty pBAD vector grown to an OD600 of 0.6 (blue, stopped after 40 min) and induced with 0.2% arabinose. All data was blanked by the fluorescence signal at 0 min, and data post-induction was smoothed by three passes though a seven-point moving average filter. The standard deviation of the baseline fluorescence in the 10 minutes prior to injection for each run was used to calculate its respective detection threshold (Table 2). Note the sharp drop in fluorescence immediately following arabinose injection, which is due to sample agitation post-injection.

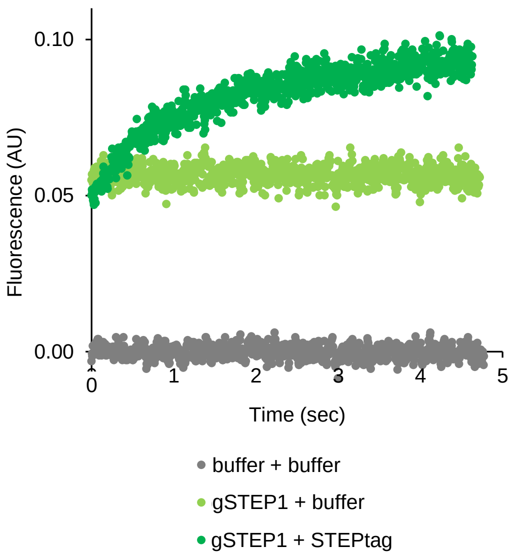


**Supplementary Figure 5. Controls for rapid-mixing stopped-flow kinetics.** All experiments were performed at 37°C in 20 mM sodium phosphate buffer containing 50 mM NaCl (pH 7.4). Fluorescence signal from the stopped-flow spectrophotometer is shown ($\lambda_{ex}$ = 485 nm, $\lambda_{em}$ = 515 nm). Injection of equal volumes of samples into the mixing chamber occurs at time = 0 sec. Contents of syringe 1 + syringe 2 are indicated. Concentrations of gSTEP1 and STEPtag were 1 µM and 5 µM, respectively.
